# Evaluation of the Ability of AlphaFold to Predict the Three-Dimensional Structures of Antibodies and Epitopes

**DOI:** 10.1101/2023.08.03.551715

**Authors:** Ksenia Polonsky, Tal Pupko, Natalia T Freund

## Abstract

Being able to accurately predict the three-dimensional structure of an antibody can facilitate fast and precise antibody characterization and epitope prediction, with important diagnostic and clinical implications. In the current study, we evaluate the ability of AlphaFold to predict the structures of 222 recently published, non-redundant, high resolution Fab heavy and light chain structures of antibodies from different species (human, *Macaca mulatta*, mouse, rabbit, rat) directed against different antigens. Our analysis reveals that while the overall prediction quality of antibody chains is in line with the results available in CASP14, other antibody regions like the complementarity-determining regions (CDRs) of the heavy chain, which are prone to higher genetic variation, generate a less accurate prediction. Moreover, we discovered that AlphaFold often mis-predicts the bending angles between the variable and constant domains within a Fab. To evaluate the ability of AlphaFold to model antibody:antigen interactions based only on sequence, we used AlphaFold-multimer in combination with ZDOCK docking to predict the structures of 26 known antibody:antigen complexes. ZDOCK succeeded in predicting 11, and AlphaFold only two, out of 26 models with medium or high accuracy, with significant deviations in the docking contacts predicted in the rest of the molecules. In summary, our study provides important information about the abilities and limitations of using AlphaFold to predict antibody:antigen interactions and suggests areas for possible improvement.

**Key Points:**

- AlphaFold was used to predict 222 new 3D hi-res atomic structures of Ab chains.
- Low accuracy was observed in the prediction of HC-CDR3 and the elbow angles.
- Predicting Ab-Ag complexes and epitope mapping using AlphaFold-Multimer was limited.

## Introduction

Antibodies (Abs) are the basis of all approved vaccines and are major correlates of protection in all vertebrates (1–4). Physiologically, antibodies are produced by B cells following immunization or infection (5). Importantly, these B cells have the ability to improve, and affinity mature their presented antibodies, while also differentiating into antibody secreting plasma cells and a specific subset of memory B cells. Subsequently, memory B cells and the antibodies they produce are largely responsible for preventing re-infection and reducing the severity of the disease during secondary encounters (6). Their exquisite specificity and affinity make antibodies an appealing class of drugs that are widely used in the clinic for the treatment of cancer (7), autoimmune disorders (8), and more recently, for infectious diseases (5). Over the last two decades, innovative engineering and single-cell and high throughput cloning techniques have significantly advanced the ability to generate new antibodies against various specific targets. Technical advances and optimized expression protocols now enable rapid generation of a large number of antibodies, so that sometimes the antibodies can be tested for their therapeutic activity in animal models within 1-2 weeks of collection of the original specimen (9). Such antibodies can be also used for diagnosis and as guides for vaccine design (10, 11).

Despite a deserved sense of achievement and accomplishment, one significant bottleneck remains within “the antibody pipeline” of every pre-clinical and clinical antibody, namely deciphering the mechanism of action. This usually involves investigation of the antibody structure and identification of the antibody binding site, i.e., the precise epitope on the target (12). Precise and detailed information regarding the epitope is crucial for understanding the antibody’s functions and for predicting possible escape mechanisms (13–15). If the 3-dimensional (3D) structure of the antibody:antigen complex is not available, there are a variety of computational tools that can be used to dock the 3D structure of the antibody with the 3D structure of antigen. Such docking algorithms have been used with various degrees of success (16, 17), but generally require separately solved crystal structures of the antibody and the antigen. Unfortunately, while the antigen structure is usually known and resolved, the atomic coordinates of the antibodies generated against it are often lacking, and their solution requires weeks, and sometimes months. Thus, despite the advances in antibody isolation and sequencing, and the new computational and structural epitope-mapping methods, delineating the structure of both the antibody and its epitope is still a major challenge (12, 18–20). Being able to model the structure of a new antibody based solely on the primary amino-acid sequence will provide a better understanding of antibody function and antibody:antigen interactions and will greatly contribute towards the use of antibodies in research and clinical applications (21–23).

Predicting protein structure based solely on the coding primary amino-acid sequences is often referred to as “the protein folding problem”. This refers to the challenge of addressing protein complexity, and the distinct quaternary structure with a significant degree of flexibility in some areas and rigidity in others (15, 24). Critical Assessment of Protein Structure Prediction (CASP) is a biennial community experiment designed to determine the state of the art in modeling protein structure (25). Participants are provided with the amino-acid sequences of target proteins and build models of the corresponding 3D structures. DeepMind’s AlphaFold AI system (26) has demonstrated remarkable results in CASP14 (25), achieving a median global distance test score of 0.92, where the global distance test score values range from 0 to a perfect score of 1.0. This was a significant leap compared to the previous year, when the score was 0.59 (27–29). However, while the analysis included 84 experimental models and 152 different protein prediction targets related to these models, it did not specifically focus on antibody evaluation, and included only two prediction targets related to a single antibody model (PDB ID 6VN1) (30). Therefore, while AlphaFold is an outstanding tool for predicting protein folding, its accuracy in modeling antibody structures remains unknown.

Antibodies share common conserved areas, whose structure is easy to predict by homology. In contrast, predicting the structure of the antibody variable regions is substantially more challenging, because their generation via a genetic recombination between randomly selected genome-encoded V, D, and J regions introduces a high degree of diversity. In addition, the sequence, and subsequently the structural diversity, are further increased by the insertion of n-p nucleotides, and later random mutations, into the coding sequence during the affinity maturation that follows antigen exposure (13, 31–33). This variability greatly reduces the availability of closely related examples from which structure prediction algorithms can learn, and thus these regions are predicted with lower accuracy, leading to a poor overall prediction accuracy for antibody structures (34, 35).

This study was designed to evaluate the prediction accuracy of AlphaFold as related to antibodies. Specifically, we asked the following questions: (1) What is the average accuracy of AlphaFold in predicting atomic structures from primary amino acid antibody sequences? (2) Which antibody chains are predicted with better or worse accuracy? (3) What structural elements within the antibody chain are accurately inferred and which regions are prone to mistakes? (4) What types of mistakes can be expected when AlphaFold is used to predict the 3D structures of antibodies? (5) How well can AlphaFold predict specific epitopes on the surface of the corresponding targets, and how does the level of accuracy compare to that of ZDOCK, which is an alternative prediction algorithm based on docking (36).

## Methods

### Data

AlphaFold prediction quality was evaluated on antibodies with crystal structures available in the Protein Data Bank (PDB) (37). A non-redundant set of antibody structures was extracted from the SAbDab antibodies database (38), with a maximum sequence similarity of 80% and a resolution cutoff of 3Å. Only structures that were released after the AlphaFold training cut-off date of April 30, 2018, were considered. Importantly, we focused on the fragment antigen-binding (Fab) regions of the antibodies, as these regions confer most of the specificity of an antibody to the corresponding antigen. Heavy and light chains were analyzed separately, and the dataset included different species (Table 1). In total, we analyzed 222 Fab chains, 95 full Fab structures, and 26 Ab-Ag complexes. Full data used in the analysis can be found at https://github.com/XseniaP/AF_evaluation/blob/master/Supplementary_Tables.xlsx.

**Table 1.**
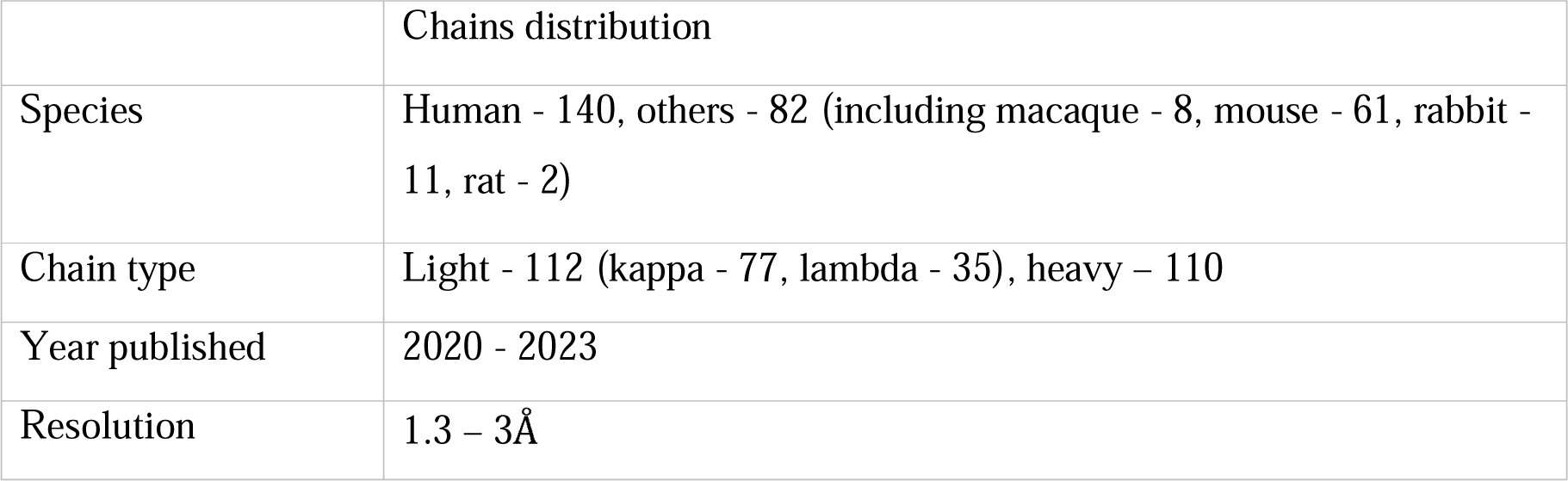
Distribution of the 222 analyzed Fab chains. High-resolution immunoglobulin Fab fragment chains were chosen for the analysis to ensure that the distance between predicted model and reference molecule is not affected by resolution. All the structures analyzed were published after AlphaFold v2.2.0. training cut-off date of April 30, 2018.

### AlphaFold

The full version of AlphaFold v.2.2.0. (https://github.com/deepmind/alphafold/) without “templates” (no homologous structure search) was used for structure predictions (39). AlphaFold also uses the BFD database, which is currently one of the largest publicly available sets of protein sequences (26, 40, 41), with the original version of this database including more than two and a half billion protein sequences clustered in families. DeepMind has already added many of the molecules predicted by the complete AlphaFold system to the AlphaFold Protein Structure Database (42), but not all structures of interest are present in the database as a whole, and we therefore decided to run AlphaFold v.2.2.0. locally in order to predict all possible structures.

### Source code and program

We developed a Python script (the Script) (https://github.com/XseniaP/AF_evaluation) to perform the calculations and combine all accuracy evaluation steps into a single pipeline (Fig. 1). The script is used to call the externally developed open-source products for alignment, superimposition, metrics calculation, and visualization, and then parses the results and moves the relevant information to the next step. In addition, the script provides on-demand calculations of the key metrics per domain and color coding, which are not covered by the externally developed packages.

**Figure 1:**
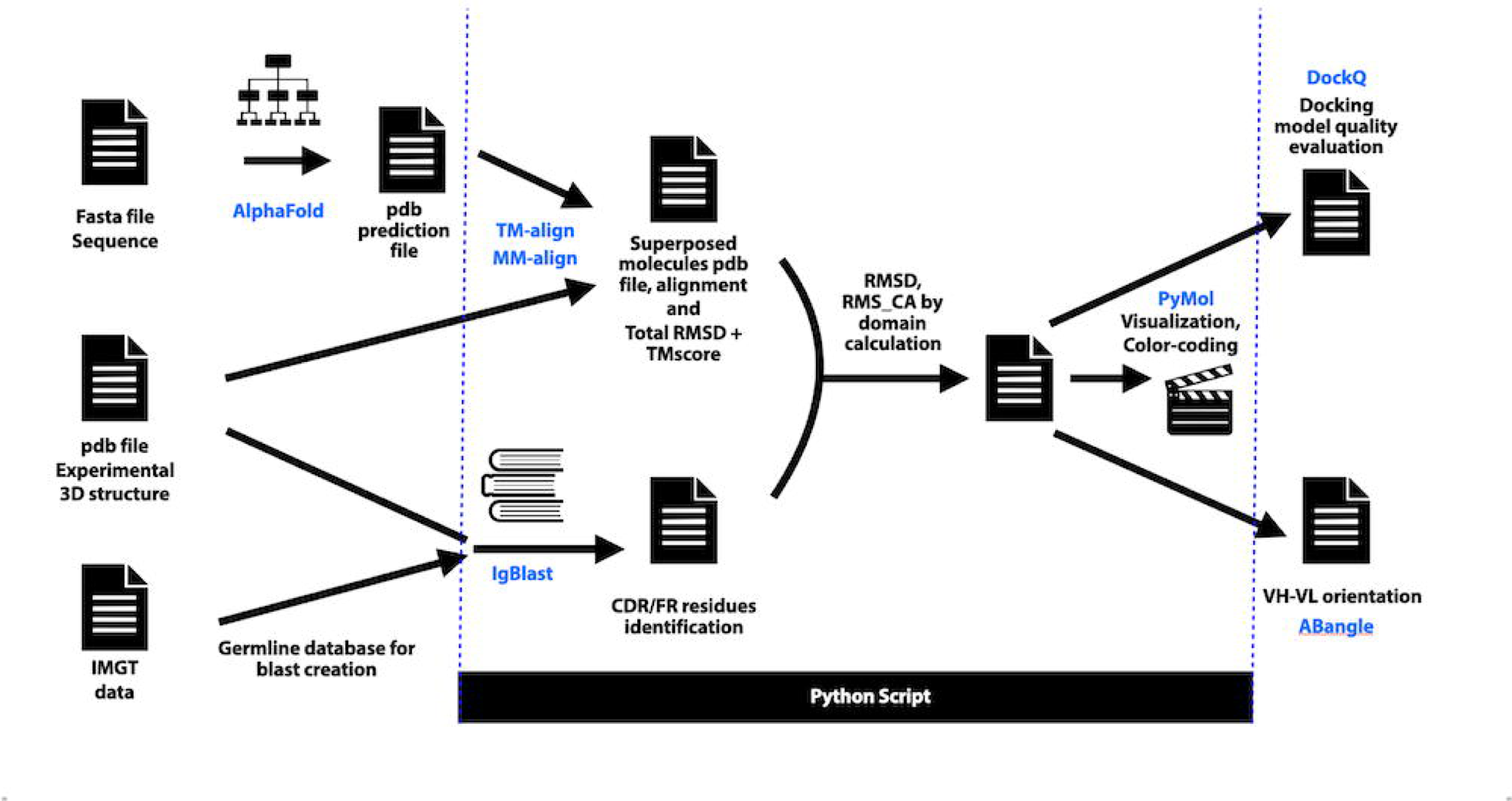
Study visualization chart. The analysis workflow starts with a target sequence in Fasta format, a pdb file of the native structure, and a pdb file of the structure predicted by AlphaFold v2.2.0, which corresponds to the input sequence. IMGT reference directory data are pre-downloaded, and a germline database is created accordingly. We then use TM-align and MM-align to superimpose the native and the predicted structures, which provides overall RMSD and TM-score measures. We also run PyIR wrapper of IgBLAST to detect regions in the sequence corresponding to the CDR/FR regions of the antibody. Thereafter, additional metrics per region are calculated to identify the regions with the lowest prediction quality. Elbow angle is calculated using RBOW software and VH-VL orientation is obtained using the ABangle computational tool. Docking model quality was evaluated by using DockQ software. We also produce PyMol scripts to color-code the molecule residues based on their prediction quality reflected by RMSD, and used ChimeraX to compare the predicted and native multimer structures.

### Identifying structural subdomains

The variable fragment of the Fab domain of the antibody is formed by the heavy and light chains and is responsible for the specificity of the antibody for its antigen. As the first step and to identify problem areas, the quality of tertiary structure predictions for 222 Fab structures was analyzed, with each chain considered separately. Next, we assessed the quality of the prediction for the structural domains of the Fab, i.e., variable versus constant domains. Lastly, we assessed the structural subdomains within each domain, i.e., the three hypervariable loops, known as complementarity determining regions (CDRs) of the heavy and the light chain (CDRH1, CDRH2, CDRH3, CDRL1, CDRL2, and CDRL3), which form the specific antigen recognition site on the surface of the antibody (43), as well as the less variable framework (FR) regions.

Kabat (44) and Chothia (45, 46) are two variable domain numbering schemes that are often used to define the location of the variable fragment regions in the sequence. Two common programs for processing immune repertoire sequencing data are IMGT (47) and NCBI IgBLAST (48), where the latter has an offline version, and Python library wrappers exist for data processing. In this study, we used the IMGT reference directory (49, 50) for immunoglobulins as a basis for generating a germline database for each species. The databases were also used as a prerequisite for the PyIR wrapper (51) in order to incorporate the IgBLAST-based mechanism into our Python script and identify the chain type as well as the CDR1/2/3, FR1/2/3 regions of the variable domain. If the location of the end of the CDR3 region was missing in the PyIR output, the result was verified and completed using known CDR3 motifs (43, 52–54) for each of the corresponding chain types, as presented in Table S3. For example, the C terminal of a heavy chain frequently juxtaposes a Trp-Gly-XXX-Gly motif and the TGGGG motif often appears as an end site of the CDR3 domain in the heavy chain. The motifs for each of the chain types were only used to verify and complement the CDR3 region in the corresponding chains when this information was missing in the output of PyIR.

### Superimposing the molecules

TM-align (55), which employs the coordinates of the backbone carbon alpha (*C_α_*) of a given protein The predicted and experimentally determined 3D structures (the native structure) were aligned by structure for superimposition and distance calculations. TM-align was run in two different modes. First, metrics were calculated using sequence-independent alignment, i.e., the superimposition was based on structural similarity. Such an analysis should produce the closest superimposition with the lowest distance between two molecules but may not align the same domains of the native and predicted molecules and thereby prevents the analysis of the prediction quality by domain or identification of the exact location at which the prediction accuracy deteriorates. We therefore repeated the calculations with sequence-dependent alignment, where the residue index correspondence between two structures is included as a constraint. MM-align (56) developed for sequence-independent alignment of complex protein-structures was used to compare predicted and native multimer structures, such as an entire Fab fragment and antibody-antigen (Ab-Ag) structures. We also used the ChimeraX MatchMaker (57) tool, which is part of UCSF Chimera (58). This algorithm allows us to superimpose complex protein structures by first creating a pairwise sequence alignment between the selected chains, and then fitting the aligned residue pairs and calculating the distance between them.

### Accuracy evaluation

CASP and Antibody Modeling Assessment (AMA) (39), where *d_i_* is defined as the distance between The prediction accuracy was calculated from parameters selected from the official metrics used by the, *i*^th^ pair of aligned residues in the metrics described below.

1. RMSD - root-mean-square deviation values of the subset of *C_α_* atoms that correspond to the residues from the target crystal structure after sequence-independent structural superimposition of the two atoms. The TM-align sequence-independent mode was applied, i.e., the superimposition was generated based on minimum distance without any constraints on the sequence alignment. RMSD values were obtained as a part of the standard TM-align output based on the following formula:

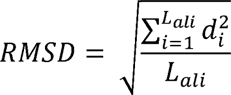

where *L_ali_* represents the number of aligned residues. MM-align was used to calculate the RMSD values based on the differences between native and predicted complex structures.
2. RMS_CA - root-mean-square deviation for the entire target structure was calculated on *C_α_* atoms based on sequence-dependent structural superimposition, which assumes residue index correspondence between the molecules. The TM-align sequence-dependent model was applied and the metrics were calculated by our script based on the following formula:

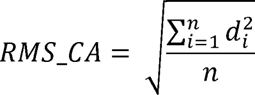

where *n* represents the total number of residues in the molecule. ChimeraX MatchMaker was used to calculate RMS_CA values based on the differences between native and predicted complex structures while creating a pairwise sequence alignment.
3. GDT – global distance test quantifies protein dissimilarity. Notably, GDT is less sensitive to outlier regions that might result from the poor prediction of a specific region (*e.g.*, a CDR loop), while the rest of the model is reasonably accurate (59). The two GDT measures are the global distance test total score (GDT_TS_) and the global distance test high accuracy score (GDT_HA_). These measures were calculated on two structurally superimposed molecules, both in sequence-dependent and sequence-independent mode. The GDT_TS_ was calculated according to the following formula:

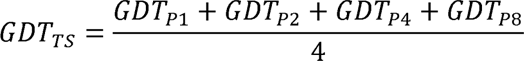 GDT_HA_ was calculated according to the following formula:

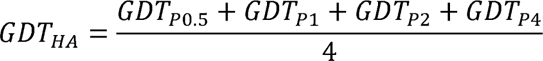

where *GDT_Pn_* denotes percentage of residues with a distance equal to or less than n <u>Å.</u> We have implemented these computations with our Python program. The formulas are those used by CASP to evaluate prediction accuracy (25, 59). The models submitted to the CASP competition are ranked based on the GDT_TS_ score, which is used as the major assessment criterion. GDT_TS_ provides an estimate of the percentage of residues predicted under specific cutoff distances, and GDT_HA_ provides an estimate of the percentage of residues predicted with high accuracy.
4. TM-score – the template modeling score is a metric for assessing the topologic similarity between two protein structures. We use the TM-score, which is provided as a standard output of the TM-align, and is defined as (55):

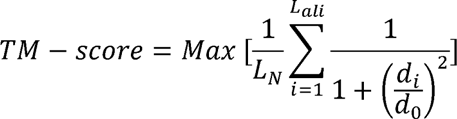

such that *L_N_* is the length of the native protein that other structures are aligned to; *L_ali_* is the number of aligned residues; 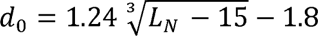 a scale to normalize the match difference and to rule out protein size dependence. This is based on an estimation of the average structure distance between aligned residues of the random related structures of the length *L_N_*. TM-score value ranges from 0 to 1, which indicates a perfect match between two structures. Scores below 0.2 indicate randomly chosen unrelated proteins. AlphaFold-Multimer (60, 61) was used to predict structures of heavy and light chains bound together. Using this analysis, we could better evaluate the prediction accuracy of AlphaFold, with the following metrics:
5. Elbow angle – a measure of the orientation between the variable and the constant domains in the Fab region. The elbow angle has been shown to increase Fab flexibility and enhance the ability of the antibody to recognize different antigens (62). Antigen binding causes an apparent shift of the angle, which reflects the conformational changes that occur (63). The elbow angle also plays an essential role in antibody assembly (24, 64). Hence, the elbow angle is critical for antibody structure functionality and an essential parameter in modeling and antibody engineering. The Fab elbow angle was computed using RBOW (62). The difference in elbow angle between the predicted and native structures was used to quantify the accuracy.
6. VH-VL orientation in Fab region of antibodies was measured by the ABangle computational tool using five angles (HL, HC1, LC1, HC2 and LC2) and a distance (dc) (65).
7. DockQ score – is a quality measure for protein-protein docking models derived by combining *F_nat_*, LRMS, and iRMS to a single score in the range [0, 1]. DockQ values can be interpreted as follows: 0.00 ≤ *DockQ* < 0.23 corresponds to “incorrect”, 0.23 ≤ *DockQ* <0.49 to “acceptable quality”, 0.49 ≤ *DockQ* < 0.80 to “medium quality”, and *DockQ* ≥ 0.80 to a “high quality” docking model (66).

### Analyzing VDJ mutations

We obtained the germline information for each of the antibody chains from the PyIR wrapper (51) output and counted the number of amino-acid substitutions in the VDJ region compared to the germline. This number was used to define four levels: low (under 5 amino acid substitutions per chain), medium (under 10), high (under 20), and extensive (20 or more).

### Visualizing and color-coding results

A PyMol v.2.5.2 (67) molecular visualization system script was written to color code the chains. ChimeraX (58) was used to visualize superimposed multimer structures.

### Docking

The Ab-Ag complex structure can be predicted by AlphaFold-Multimer. It can also be predicted by molecular docking, which takes known or predicted 3D structures of an antibody and its corresponding antigen as input and returns a single 3D structure of the Ab-Ag complex. Docking was performed using the ZDOCK 3.0.2 web server (36).

## Results

### Overall prediction accuracy

AlphaFold predictions of the published structures of 222 antibody chains, generated scores that were generally similar to the official CASP14 results for protein prediction (27–29, 68): The mean and median RMSD scores were 2.38 Å and 2.13 Å, respectively with 43.69% of the chains scoring lower than 2 Å, and 70.27% of the sample scoring lower than 3 Å (Table 2). The average *GDT_TS_* and *GDT_HA_* values for all the predictions in our set were 0.72 and 0.50, respectively, where a *GDT_TS_* of 0.72 (out of 1.0) reflects a good approximation to the native 3D structures. The *GDT_HA_* score, which penalizes larger deviations from the native structure, suggests that, on average, 50% of the positions within a Fab can be predicted with very high accuracy. The average TM-score for all individual heavy and light chains in our analysis was 0.83, which represents not only the same fold but actually a very close match between the predicted and native structures.

**Table 2.**
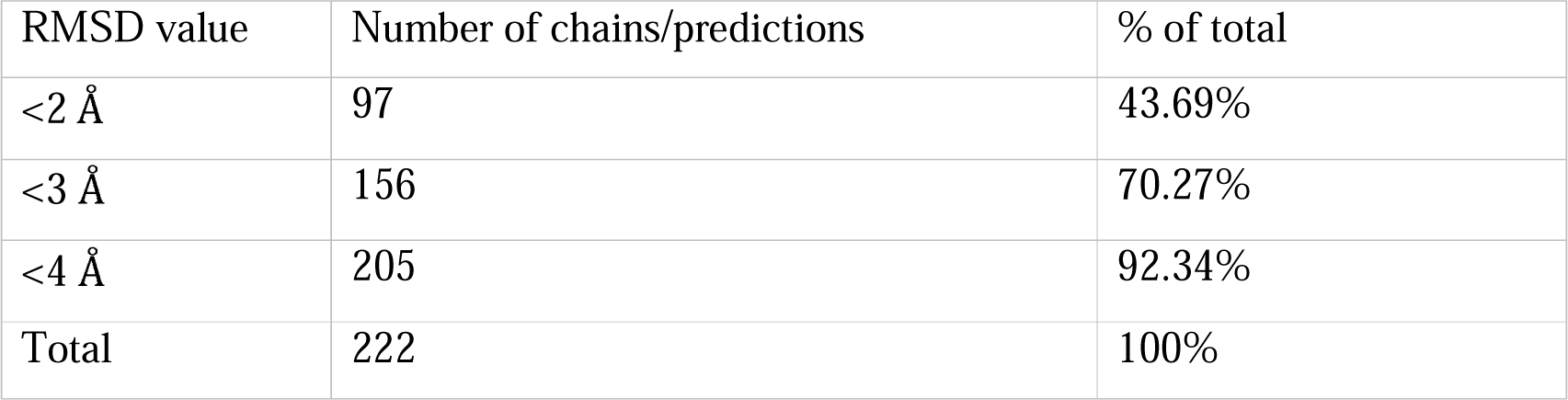
222 predicted chains’ distribution by RMSD value against corresponding original molecules.

| RMSD value | Number of chains/predictions | % of total |
| --- | --- | --- |
| <2 Å | 97 | 43.69% |
| <3 Å | 156 | 70.27% |
| <4 Å | 205 | 92.34% |
| Total | 222 | 100% |

### Prediction accuracy for difference chain types and for different species

The mean RMSD values were higher in the heavy chains than the light chain: 2.68 Å compared to 2.09 Å, respectively (two-tailed t-test: p-value 1.98 x10^−5^, Table 3). The highest observed values excluding outliers according to IQR method were 4.96 Å for heavy chains *vs.* 4.81 Å for light chains, while the lowest values excluding outliers according to IQR method were 1.05 Å for heavy chains *vs.* 0.72 Å for light chains (Fig. 2a). Lambda light chains had a higher average RMSD value (two-tailed p-value 2.12 x 10^−5^) than kappa light chains (2.74 vs 1.79 Å respectively, Table 3, Fig. 2b). Similar results of 0.72 and 0.77 for lambda and kappa light chains, respectively were obtained when the *GDT_TS_* score was used to quantify prediction accuracy (Supplementary Fig. S1). Comparing Fab *Macaca mulatta*, with a mean RMSD value of 3.43 Å compared to other species for which the mean chains from different species revealed a slight, albeit statistically significant, increase in RMSD in RMSD was in the range 1.61 - 2.52 Å (ANOVA one-way test p-value 4.13 x 10^−4^, Supplementary Fig. S2). The lower predictability of *Macaca mulatta* Fabs may stem from the underrepresentation of *Macaca mulatta* structures in the training set of the AlphaFold v2.2.0. model as well as the low representation in our set.

**Figure 2:**
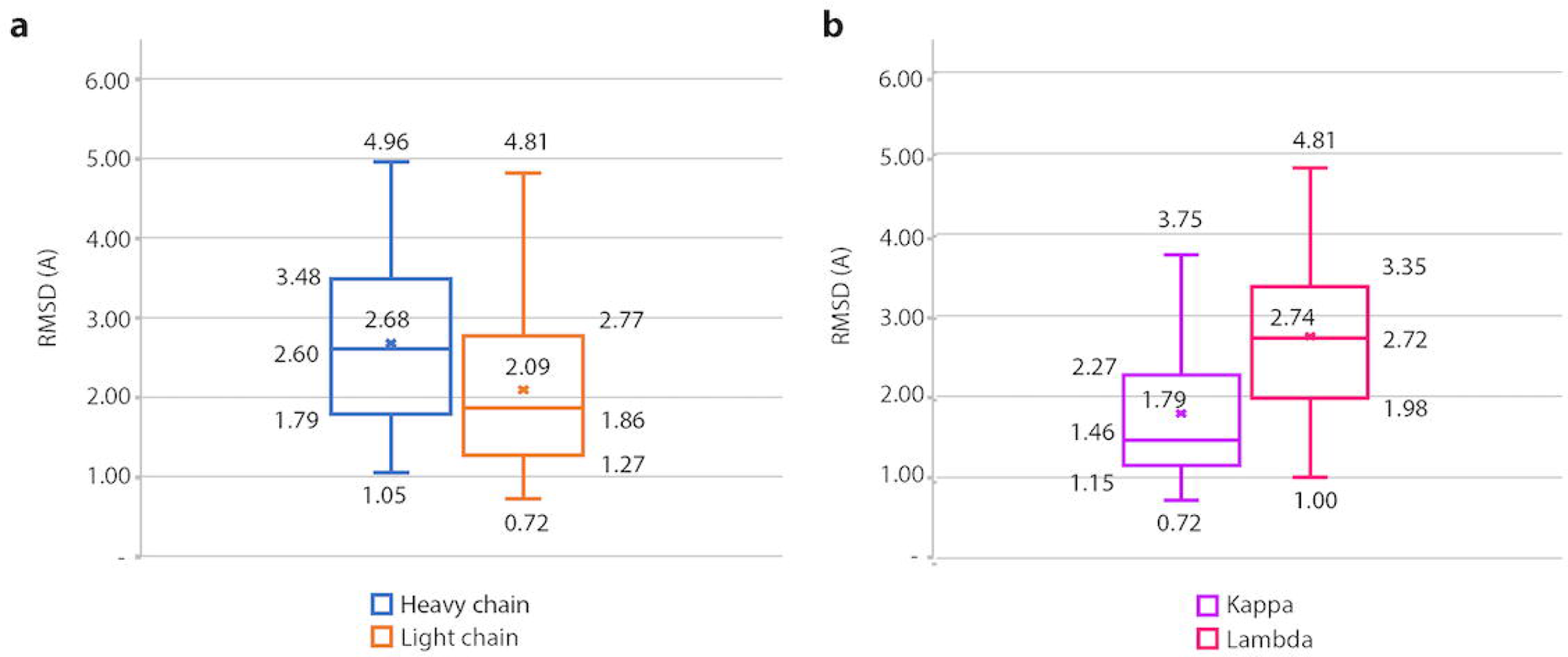
Prediction quality evaluation by chain type and light chain class. Boxplot of RMSD (Å) data presenting the distance between superimposed native and predicted molecules for (a) light and heavy chains and (b) kappa and lambda chains.

**Table 3.** Predicted chains’ distribution by RMSD value against corresponding native molecules according to chain type and light chain class.

|  | Heavy chains |  | Light chains |  | Light kappa chains |  | Light lambda chains |  |
| --- | --- | --- | --- | --- | --- | --- | --- | --- |
| RMSD | Number of predictions | % of total | Number of predictions | % of total | Number of predictions | % of total | Number of predictions | % of total |
| <2 Å | 36 | 32.73% | 61 | 54.46% | 52 | 67.53% | 9 | 25.71% |
| <3 Å | 65 | 59.09% | 91 | 81.25% | 68 | 88.31% | 23 | 65.71% |
| <4 Å | 96 | 82.27% | 109 | 97.32% | 77 | 100.00% | 32 | 91.43% |
| Total | 110 | 100% | 112 | 100% | 77 | 100% | 35 | 100% |
| Mean value | 2.68 |  | 2.09 |  | 1.79 |  | 2.74 |  |

### The number of somatic hypermutations within the VDJ region has little effect on the AlphaFold prediction accuracy

As the next step, we tested the hypothesis that prediction accuracy is affected by the number of VDJ mutations. To this end, we grouped the analyzed antibody chains by the number of VDJ amino acid substitutions from the germline into low (< 5), medium (between 5 and 9), high (between 10 and 19), and 2.56 Å, respectively. Although the differences among the groups are statistically insignificant (p- and extensive (20 or more). The resultant mean RMSD values for each group were 2.00, 2.47, 2.42, value > 0.05 in ANOVA), the value for the low group is significantly lower than that for all the others (p-value < 0.05 in a one-tailed t-test), even though the difference is represented by a small value (below 0.5 Å).

### Difficulty in predicting the correct angles within the antibody

The chains with a lower prediction quality and higher RMSD were examined more closely. As a case study, we selected the heavy chain fragment from the human anti-HIV-1 neutralizing antibody BG24 Fab (PDB: 7UCE) (69). Discrete AlphaFold predictions of the heavy chain constant and variable domains were highly accurate: RMS_CA of 0.48 Å and 0.42 Å for individually predicted constant resulted in an RMS_CA value of 4.93 Å, indicating a mediocre prediction (Fig. 3c). Superimposing and variable domains, respectively (Fig. 3a-b). However, a consideration of the entire heavy chain the individual domain predictions onto the prediction of the heavy chain (Fig. 3d) suggests that the lack of fit is not caused by individual domain predictions, but rather reflects a difficulty in predicting the angle between the variable and constant domains within the chain. This caused large deviations in one of the domains, leading to the high RMS_CA value.

**Figure 3:**
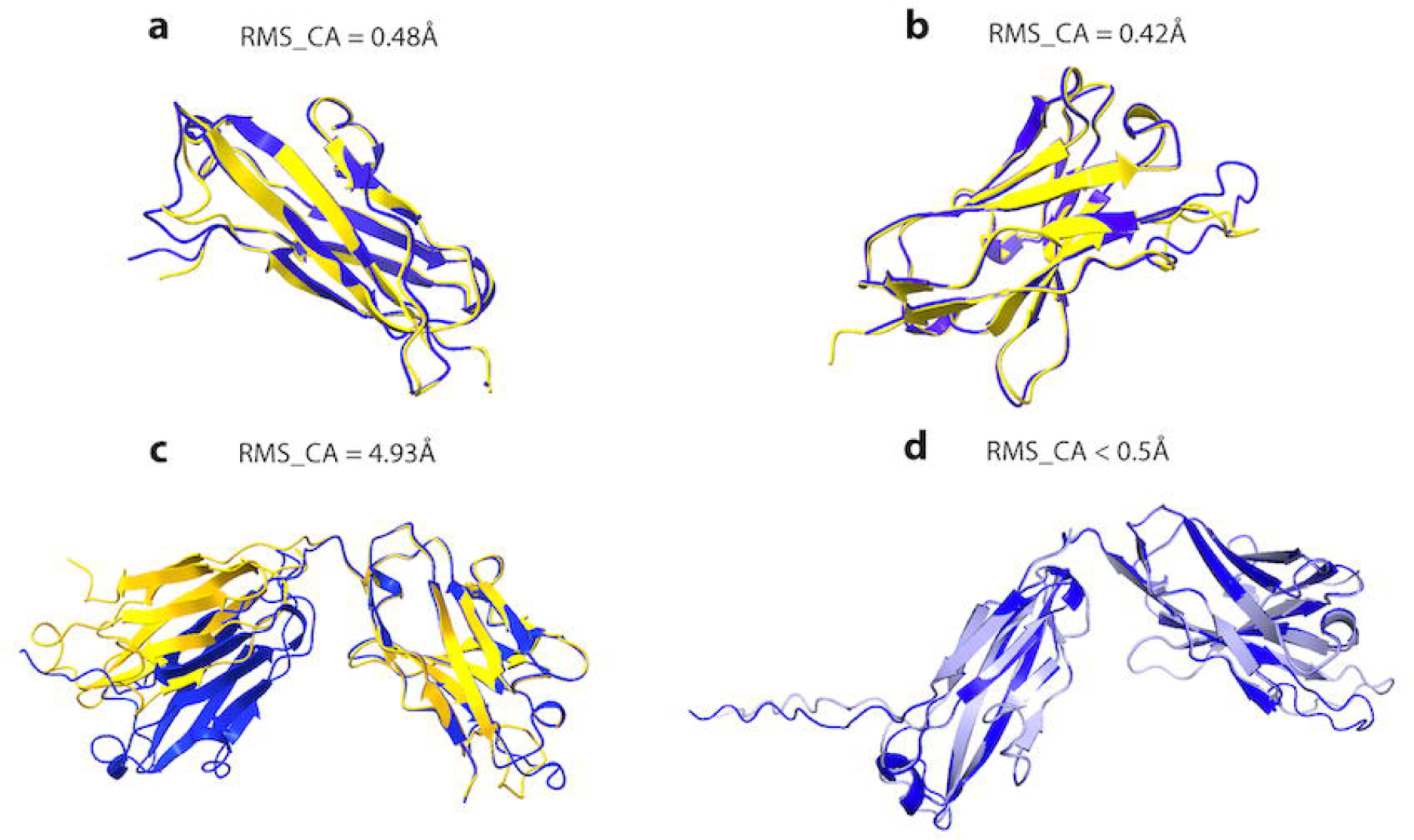
Structural analysis of the Fab fragment from the anti-HIV-1 neutralizing antibody BG24 heavy chain (PDB: 7UCE) superimposed on the AlphaFold prediction according to the sequence-dependent alignment. Cartoon diagrams of the heavy chain of the native 3D structure (in yellow) superimposed onto the same domain individually predicted by AlphaFold v2.2.0. (in blue). (a) constant domain; (b) variable domain; (c) the complete heavy chain (variable and constant domains) presented as a cartoon diagram; (d) cartoon diagrams of the variable and constant domains predicted separately (in light gray) but superimposed over the complete predicted heavy chain 3D structure (in blue).

Light and heavy chains exist as a heteromeric complex, and their correct assembly and interaction is critical for molecular function. It is therefore important to predict the entire Fab accurately. AlphaFold-Multimer predictions for 95 entire Fabs, produced RMS_CA scores ranging from 0.67 Å to 4.65 Å, with a median value of 2.14 Å. Of these Fabs, 48.42% had scores lower or equal to 2 Å However, there was a significant correlation of *R*^2^ = 0.78 between the total RMS_CA of the Fabs (Table 4). The prediction accuracy was not strongly correlated with the elbow angle itself (Fig. 4a). and the absolute value of the elbow angle deviations between the predicted and native structures (Fig. 4b). This suggests that the errors in angle estimation are responsible for the poor prediction of the Fab structure. Furthermore, our analyses indicate that AlphaFold tends to overestimate, the elbow angle: with average values of 161.0 and 172.6 degrees, respectively for the native and predicted Fabs (two-tailed t-test: p-value 0.0035). Moreover, the range of predicted elbow angles was more restricted, with a standard deviation of 30.83 and 22.32 degrees for the elbow angles of native and predicted structures, respectively. Fabs for which the predicted elbow angle deviated insignificantly from the native elbow angle (difference under 20 degrees) demonstrated high prediction quality, with a mean RMS_CA of 1.45 Å (Fig. 4c). The analysis of VH-VL orientation angles did not detect domains (Fv regions) were all predicted with a very high accuracy and average RMSD of 0.47 Å and significant differences between the native and predicted structures. In addition, ten selected variable 1.13 Å between pruned atom pairs and across all pairs respectively (Supplementary Table S1). This analysis reconfirmed our conclusions about high prediction accuracy of VH-VL orientation angles. Overall, the results demonstrate that the difficulty in predicting the angle between the variable and constant domains in the Fab region is a general issue with AlphaFold.

**Figure 4:**
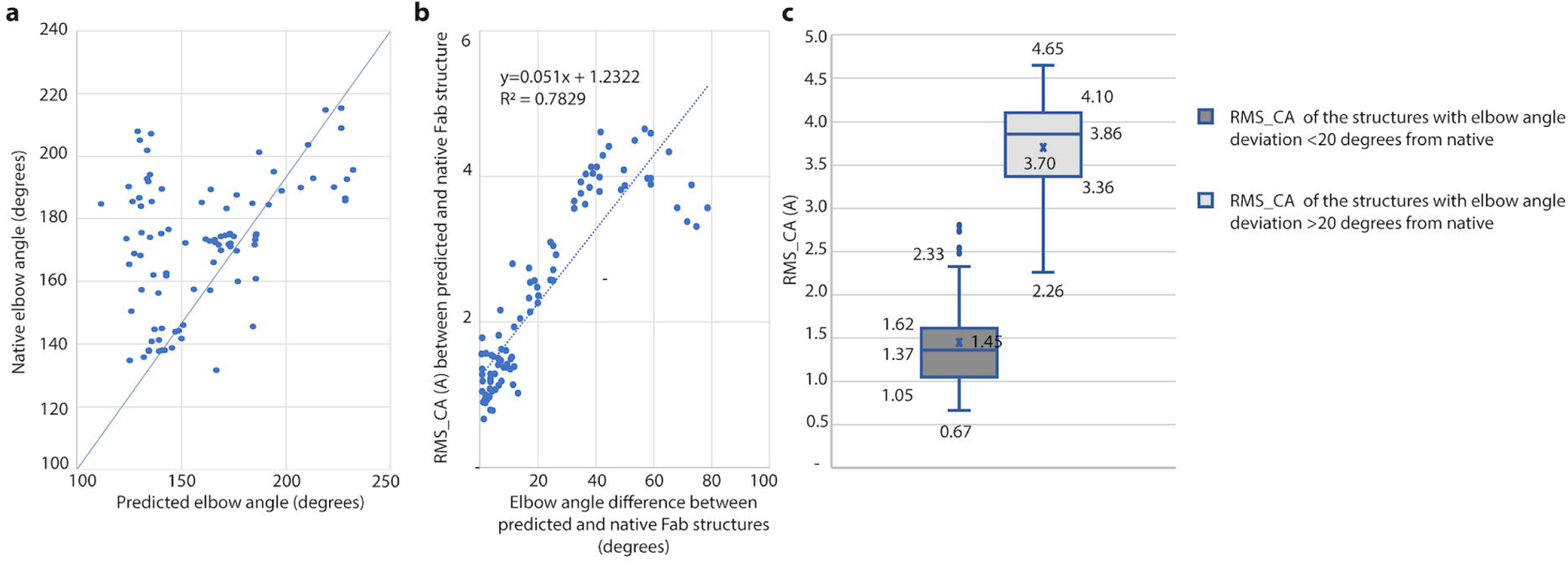
Evaluation of the elbow angle deviations correlation with prediction quality. RMS_CA (Å) in correlation with (a) scatter plot of elbow angles (degrees) data in native vs. predicted Fab structures; (b) absolute elbow angle difference in degrees between predicted and native Fabs; (c) boxplot of RMS_CA (Å) data displaying the distance between superimposed native and predicted molecules for the Fab structures for two groups of Fabs: with elbow angle difference between native and predicted of < 20 degrees, and >= 20 degrees.

**Table 4.**
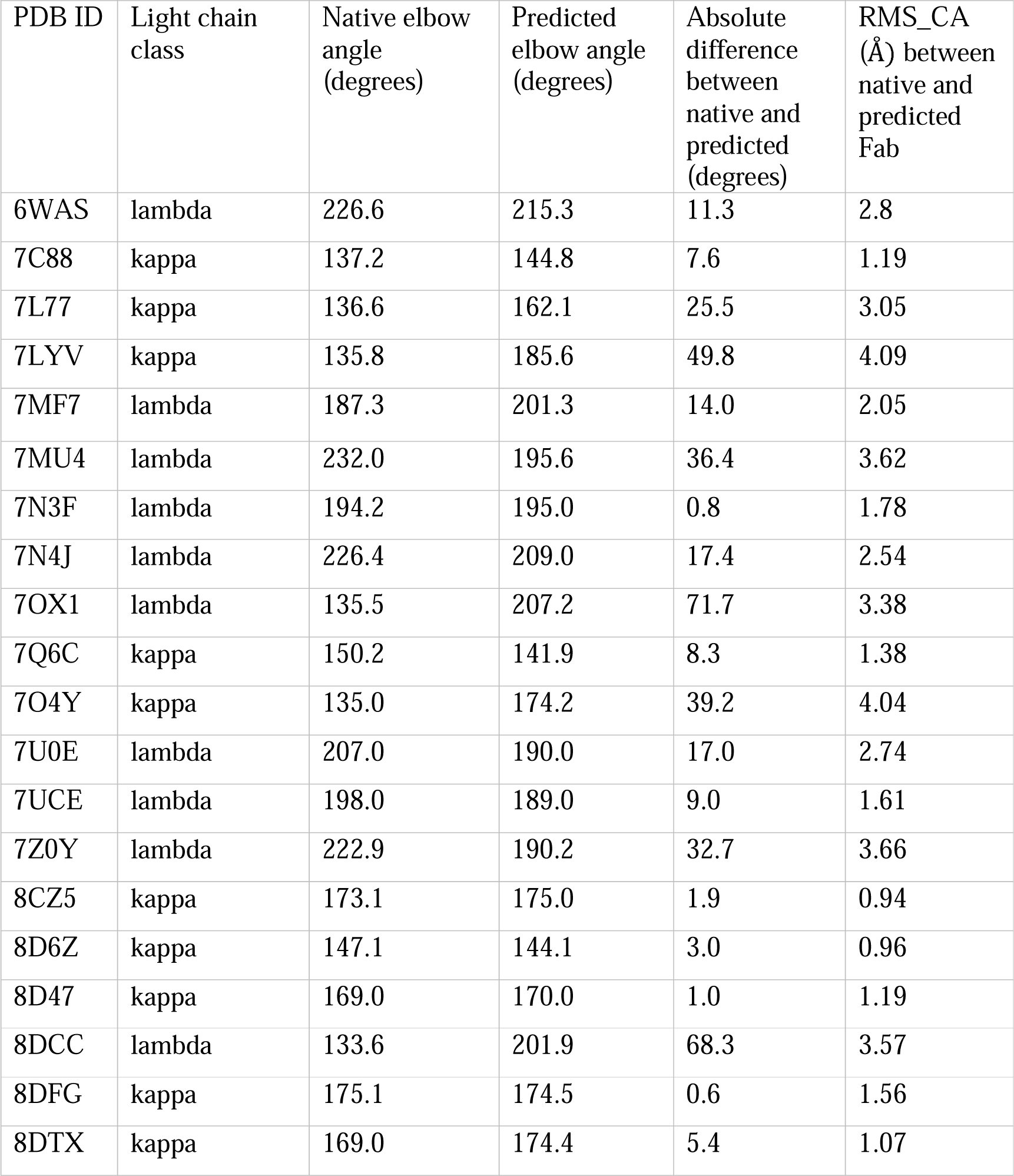
Elbow angle and RMS_CA (Å) between 20 native and predicted sequence-dependently superimposed structures of predicted and corresponding native Fab molecules (heavy and light chain combined)

| PDB ID | Light chain class | Native elbow angle (degrees) | Predicted elbow angle (degrees) | Absolute difference between native and predicted (degrees) | RMS_CA (Å) between native and predicted Fab |
| --- | --- | --- | --- | --- | --- |
| 6WAS | lambda | 226.6 | 215.3 | 11.3 | 2.8 |
| 7C88 | kappa | 137.2 | 144.8 | 7.6 | 1.19 |
| 7L77 | kappa | 136.6 | 162.1 | 25.5 | 3.05 |
| 7LYV | kappa | 135.8 | 185.6 | 49.8 | 4.09 |
| 7MF7 | lambda | 187.3 | 201.3 | 14.0 | 2.05 |
| 7MU4 | lambda | 232.0 | 195.6 | 36.4 | 3.62 |
| 7N3F | lambda | 194.2 | 195.0 | 0.8 | 1.78 |
| 7N4J | lambda | 226.4 | 209.0 | 17.4 | 2.54 |
| 7OX1 | lambda | 135.5 | 207.2 | 71.7 | 3.38 |
| 7Q6C | kappa | 150.2 | 141.9 | 8.3 | 1.38 |
| 7O4Y | kappa | 135.0 | 174.2 | 39.2 | 4.04 |
| 7U0E | lambda | 207.0 | 190.0 | 17.0 | 2.74 |
| 7UCE | lambda | 198.0 | 189.0 | 9.0 | 1.61 |
| 7Z0Y | lambda | 222.9 | 190.2 | 32.7 | 3.66 |
| 8CZ5 | kappa | 173.1 | 175.0 | 1.9 | 0.94 |
| 8D6Z | kappa | 147.1 | 144.1 | 3.0 | 0.96 |
| 8D47 | kappa | 169.0 | 170.0 | 1.0 | 1.19 |
| 8DCC | lambda | 133.6 | 201.9 | 68.3 | 3.57 |
| 8DFG | kappa | 175.1 | 174.5 | 0.6 | 1.56 |
| 8DTX | kappa | 169.0 | 174.4 | 5.4 | 1.07 |

### The accuracy of predicting different structural elements within Fabs

Our analysis of the accuracy of predicting the various domains within an antibody, indicate that CDR3 in the heavy chain and CDR1 and CDR3 in the light chain are predicted with lower accuracy: heavy chain mean RMSD values for CDR1/2/3 and FR1/2/3 were 2.50 Å, 2.24 Å, 3.60 Å, 1.99 Å, 1.81 Å, and 1.74 Å, respectively (Fig. 5a), while the light chain mean RMSD values for CDR1/2/3 and FR1/2/3 were 2.40 Å, 1.58 Å, 2.43 Å, 1.94 Å, 1.72 Å, and 1.71 Å (Fig. 5b). When considering the heavy chains: the mean RMSD values for heavy chain helices and sheets were 3.04 Å and 2.18 Å, secondary structures within the Fabs, helices were more difficult to predict than sheets in antibody respectively, with a two-tailed t-test p-value of 2.3 x 10^−6^, Fig.5c,. The corresponding mean RMSD values in the light chain were 1.87 Å and 1.76 Å, respectively with no statistically significant difference (Fig. 5d). These results were reconfirmed when the analysis was repeated by comparing sequence-dependent structural superposition: the RMS_CA value for CDR3 in heavy chains was 4.2 Å with other regions ranging between 1.53 and 2.35 Å (Fig. 5e), and the RMS_CA values for CDR1and CDR3 in light chains were 2.80 Å and 2.79 Å respectively with other regions ranging between 1.80 Å and 2.12 Å (Fig. 5f).

**Figure 5:**
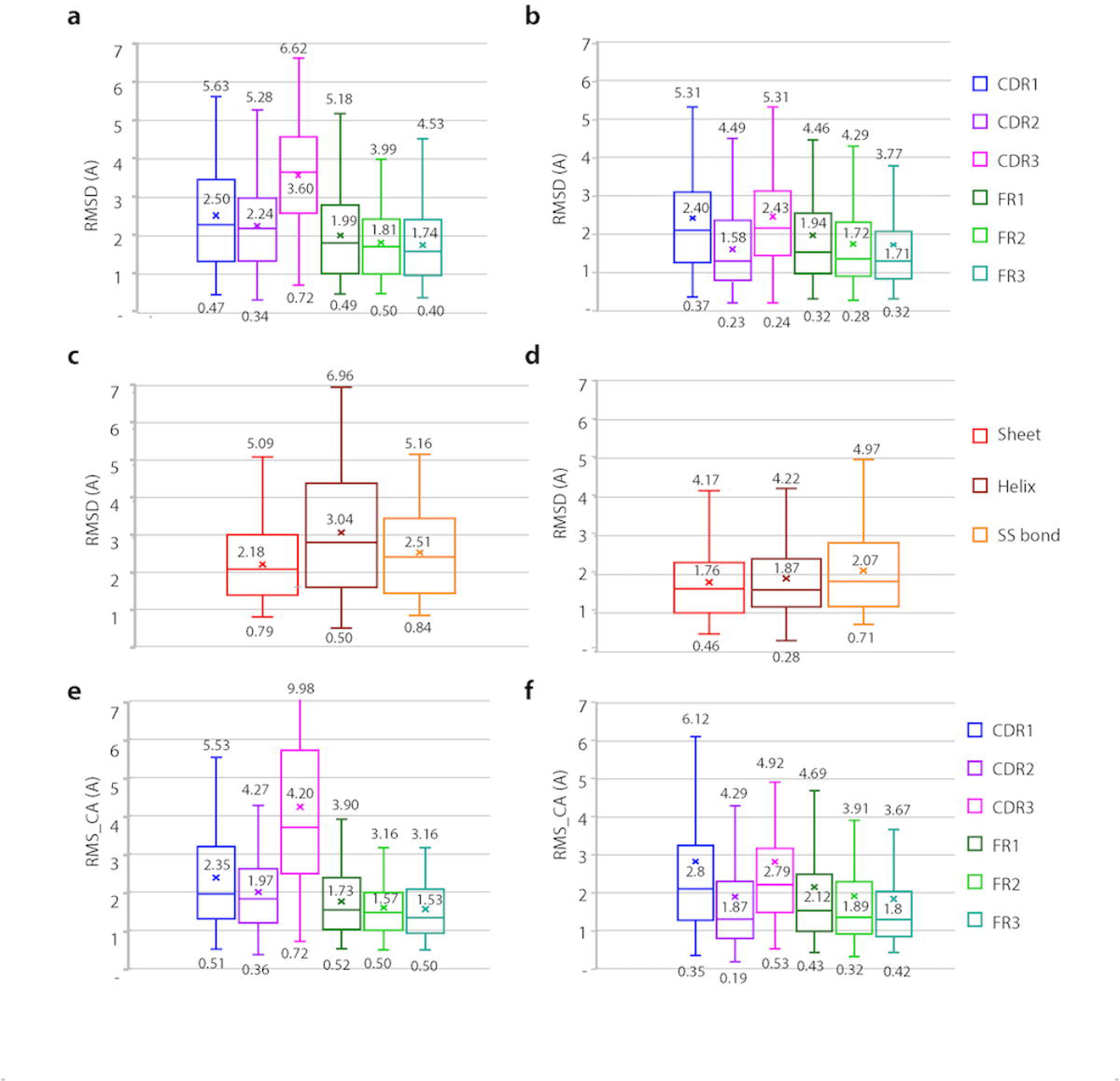
Prediction quality evaluation by subregion and by secondary structure. Boxplot of data displaying the distance between superimposed native and predicted molecules by subregion (CDR1/2/3, FR1/2/3) and by secondary structure: - RMSD (Å) by complementarity determining regions and framework regions in Fab (a) heavy chain and (b) light chain - RMSD (Å) by secondary structure in Fab (c) heavy chain and (d) light chain - RMS_CA (Å) by complementarity determining regions and framework regions in Fab (e) heavy chain and (f) light chain All superimpositions were sequence-independent in (a-d) and sequence-dependent in (e-f).

### Docking outperforms AlphaFold-Multimer in predicting accurate Ab-Ag complexes

As the next step, we tested four alternative options for predicting 26 different Ab-Ag complexes. (1) We assumed that the structure of both the antibody and the antigen are known and used the ZDOCK docking program to predict the structure of the complex. (2) First, we predicted the structure of the antibody alone using AlphaFold-Multimer, and then we predicted the docking, using ZDOCK, with the atomic coordinates of the native structure of the antigen. (3) We predicted the structure of the antigen and the antibody using AlphaFold and AlphaFold-Multimer respectively, and then docked them using ZDOCK. (4) We skipped the docking altogether and asked AlphaFold-Multimer to predict the entire Ab-Ag complex from the protein sequences of the light chain, the heavy chain and the antigen. We define docking prediction as medium quality if the DockQ score is in the range [0.49,0.8), and as high quality if the DockQ score is equal to or greater than 0.80. The results indicated that when the native structures of all components were used as input for ZDOCK (alternative 1), 11 of the 26 complexes were predicted with medium or high quality (DockQ score ≥ 0.49). In contrast, 2 of the 26 complexes predicted solely from the protein sequences by AlphaFold-Multimer (alternative 4) were of medium to high quality, with a DockQ score ≥ 0.49 (Supplementary Table S2). Significantly, none of the complexes was predicted with high accuracy when the predicted structures of both the antibody and the antigen (alternative 3), or the predicted structure of antibody and the native antigen structure (alternative 2) were used as input for ZDOCK. In general, as expected, docking with native structures as input provided a more accurate docking prediction than docking with predicted structures. Comparing docking with native antibody and antigen structures with predicting the entire complex using AlphaFold-Multimer, revealed much higher levels of accuracy with ZDOCK docking. Of note, although both complexes that AlphaFold predicted with medium to high quality also scored high for individually predicted antigen or antibody, this does not always guarantee an accurate prediction of the complex by AlphaFold-Multimer.

**Figure 6:**
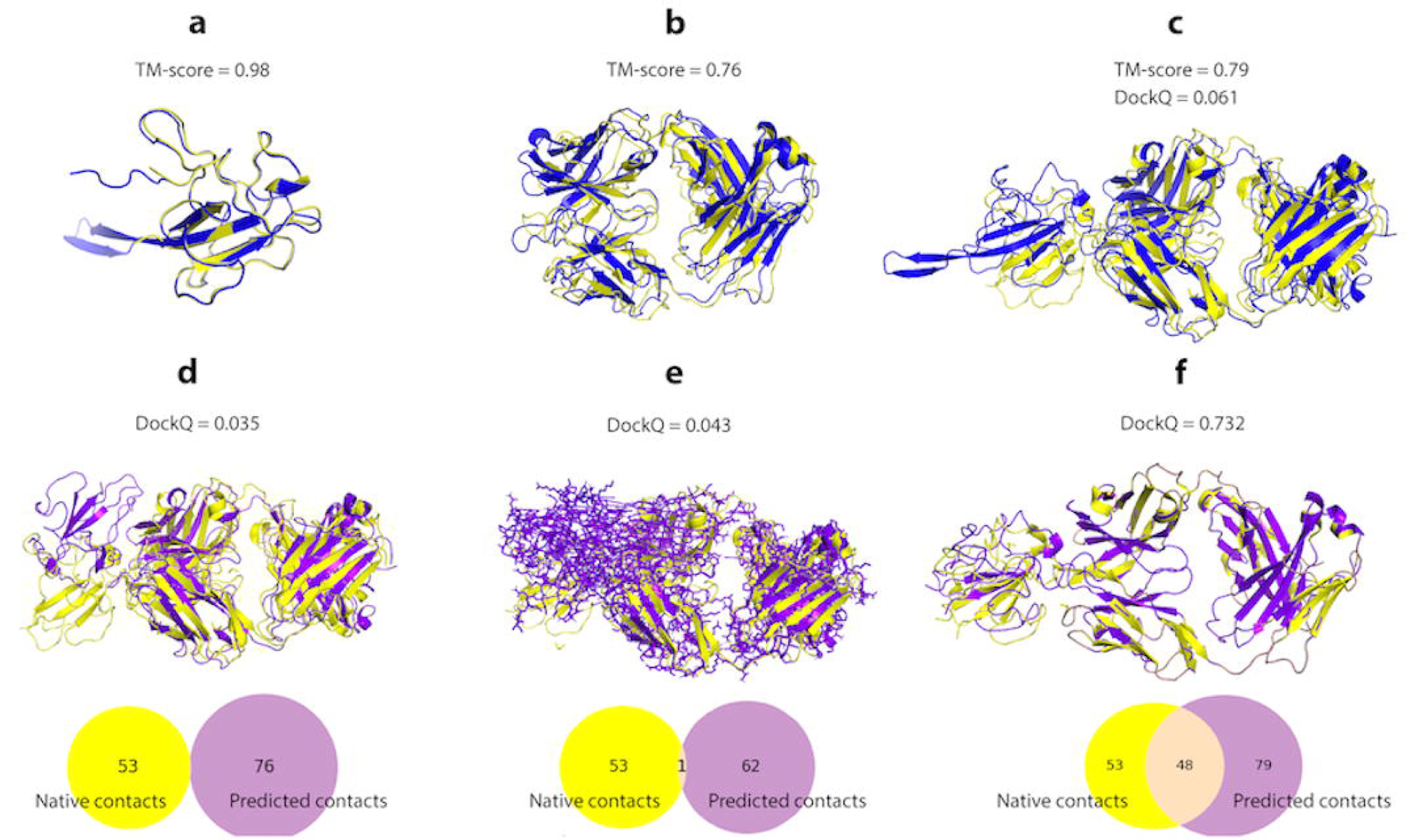
Ab-Ag multimer complex prediction and docking comparison example (PDB ID: 7N3D) (a) Native (yellow) Ag 3D structure superimposed on the AlphaFold prediction (blue); (b) Native (yellow) Ab Fab 3D structure superimposed on the AlphaFold-Multimer prediction (blue); (c) Ab-Ag complex native (yellow) 3D structure superimposed on the AlphaFold-Multimer prediction (blue); (d) Ag native structure docked by ZDOCK with Ab Fab structure predicted by AlphaFold (violet) superimposed on the native complex (yellow); (e) Ag and Ab Fab structures predicted by AlphaFold docked by ZDOCK (violet) superimposed on the native complex (yellow); (f) Ag native structure docked by ZDOCK with the Ab Fab native structure (violet) superimposed on the native complex (yellow) Venn diagrams in (d-f) show the number of docking contacts in native and predicted structures and their overlap, which represents the contacts predicted accurately.

### Prediction speed and technical requirements

One of the challenges faced while working with AlphaFold v2.2.0. was a technical GPU requirement. Given that the GPU requirements are met, it could take up to 30 minutes to predict a 216-residue long molecule. To overcome these shortcomings, we tested several alternative approaches. One approach was to utilize ColabFold (70), a web server that permits AlphaFold to run faster online (e.g., it took us up to 10 minutes to run the prediction based on a 216-residue sequence, as opposed to 30 minutes using the full version of AlphaFold v2.2.0.). In addition to ColabFold online, we also tested ABodyBuilder2 (ImmuneBuilder) (72), which is another fast deep learning model for antibody Fv region prediction. ABodyBuilder2 required about 30 seconds to predict a single antibody Fv region and the prediction accuracy for 10 tested molecules was similar to that of ColabFold online for the pruned atom pairs and somewhat better across all atom pairs (Supplementary Table S1). In October 2022, Meta AI released an alternative faster and more accessible solution as a part of its sequence-to-structure predictor ESMFold and the ESM Metagenomic Atlas database. This claimed to be 60-fold faster than state-of-the-art predictions while maintaining resolution and accuracy (71). The Atlas serves as an open database of 617 million predicted protein structures and allows a rapid prediction to be obtained from a sequence. When we applied this option to 222 Ab chains with a mean amino-acid sequence length of 221 residues, the running time was even faster, with an average of 9.3 seconds required to predict a single structure (Table 5). Moreover, more than 85% of the tested Ab sequences were predicted in under 13 seconds (Table 6). When we evaluated the prediction quality of these test molecules compared to AlphaFold predictions, the mean value of RMSD was 2.53 Å for ESMFold and 2.38 Å for AlphaFold. These predictions fall into the RMSD range of 1-2 Å, while the AlphaFold results had a higher number of differences in accuracy are not statistically significant (Fig. 7). For both algorithms, most of the predictions in the range under 2 Å (Fig. 7b). Overall, we can conclude that the prediction quality of ESMFold is very similar to that of AlphaFold, and that the shorter time required by ESMFold provides an accessibility advantage.

**Figure 7:**
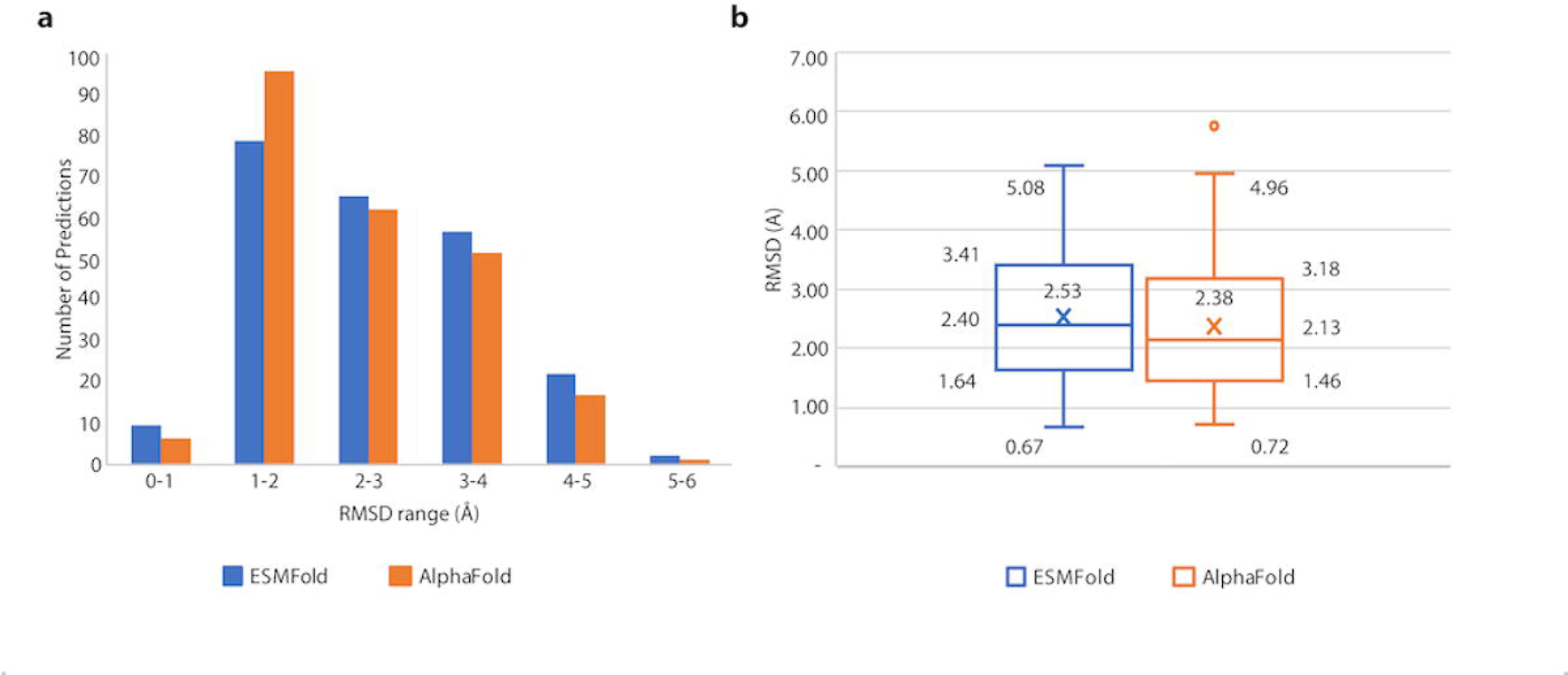
Distribution of 222 predictions by ESMFold and AlphaFold arranged by prediction quality, RMSD (Å) **(a)** Column chart of the prediction distribution by RMSD range (Å); **(b)** Box plot of RMSD (Å) data for 222 different predictions by ESMFold and AlphaFold;

**Table 5.** 20 chains prediction quality comparison between ESMFold and AlphaFold: RMSD (Å) between superimposed predicted and native molecule.

| PDB_ID & chain | ESMFold Prediction time, sec | ESMFold RMSD, Å | ESMFold TM-score | AlphaFold, RMSD, Å | AlphaFold TM-score |
| --- | --- | --- | --- | --- | --- |
| 7DOH_H | 5 | 1.87 | 0.90 | 3.83 | 0.71 |
| 7DOH_L | 10 | 3.14 | 0.77 | 0.87 | 0.98 |
| 7L77_H | 6 | 3.75 | 0.73 | 2.68 | 0.82 |
| 7L77_L | 8 | 0.91 | 0.98 | 2.01 | 0.89 |
| 7O31_H | 10 | 3.17 | 0.80 | 1.55 | 0.95 |
| 7O31_L | 10 | 4.58 | 0.63 | 1.90 | 0.90 |
| 7OX3_A | 7 | 2.33 | 0.86 | 2.34 | 0.86 |
| 7OX3_B | 15 | 0.83 | 0.98 | 3.08 | 0.78 |
| 7RM3_A | 9 | 2.41 | 0.86 | 1.50 | 0.94 |
| 7RM3_B | 12 | 1.29 | 0.95 | 1.85 | 0.91 |
| 8CZ5_H | 12 | 3.49 | 0.72 | 4.04 | 0.68 |
| 8CZ5_L | 6 | 2.73 | 0.81 | 1.31 | 0.95 |
| 8D47_H | 7 | 1.88 | 0.88 | 3.01 | 0.83 |
| 8D47_L | 6 | 1.57 | 0.93 | 1.27 | 0.95 |
| 8DFH_H | 7 | 4.77 | 0.61 | 2.02 | 0.58 |
| 8DFH_L | 6 | 2.33 | 0.52 | 1.55 | 0.93 |
| 8DTX_H | 6 | 1.52 | 0.94 | 3.28 | 0.75 |
| 8DTX_L | 6 | 2.49 | 0.84 | 1.20 | 0.97 |
| 8C3V_H | 21 | 2.24 | 0.89 | 1.54 | 0.94 |
| 8C3V_L | 5 | 0.89 | 0.97 | 2.88 | 0.79 |

**Table 6.** ESMFold 222 Ab chains prediction time distribution.

| Prediction time (sec) | Number of predictions | % of total |
| --- | --- | --- |
| 5-7 | 95 | 42.8% |
| 8-10 | 50 | 22.5% |
| 11-13 | 44 | 19.8% |
| 14-16 | 30 | 13.5% |
| 17+ | 3 | 1.4% |
| Total | 222 | 100% |

## Discussion

Understanding the antibody structure and subsequently, the antibody:antigen interactions provide a basis for the rational design of vaccines and drugs. The golden standard for structure determination remains X-ray crystallography, which provides high resolution atomic coordinates of the protein backbone and side chains, as well as the atomic interactions between the different protein groups. However, this method requires a very high degree of expertise, tends to be expensive in terms of time and money, and is not always applicable. For these reasons, a variety of methods have been developed to predict the antibody structure from the coding sequence alone.

Computational methods for protein structure prediction from the amino-acid sequence have been available since CASP1 was launched in 1994. Earlier computational methods were mostly homology-based, but although they made it possible to predict the 3D structures of simple small proteins such as PDB 2JZ5:A, with a known template with a very high sequence similarity, they were not successful in predicting the structures of larger and more conformationally challenging molecules like antibodies (73, 74). In addition, although more advanced protein prediction methods demonstrated a considerable degree of improvement, they were still inaccurate when analyzing large protein molecules, and were therefore generally not applicable for deciphering complex molecular interactions, and even less applicable for drug or vaccine design (75–77). The advent of artificial intelligence and development of new machine-learning methods provided a significant leap in the protein structures with a median *GDT_TS_* score of 0.92. While the analysis included many complex ability to predict protein structures. In 2020, AlphaFold demonstrated an improved ability to predict and structurally challenging molecules, the prediction of antibody structures and interactions has not been thoroughly evaluated. Although a number of AlphaFold protein predictions have been reported over the past few years (78–80), these did not address the specific issues of antibodies. In addition, most of these studies lacked local quality assessment of the predictions and did not try to estimate the reliability of the different subdomains of the predicted antibody or identify the areas that were the most difficult to predict.

Here, we evaluated the ability of AlphaFold to predict antibody 3D structures from a coding linear sequence, by employing an algorithmic pipeline that compares the 3D structure of the predicted antibody chain molecule with the native X-ray crystallography structure. Importantly, the system scores the structural subdomains as well as the whole molecule. This methodology enabled a rapid comparison of 222 predicted structures with the corresponding published PDB atomic structures. The results allowed us to identify the problematic areas in the predicted molecules which can then be further analyzed and used to refine the models. We also predicted the 222 antibody chains by ESMFold released by Meta AI and compared the performance with the AlphaFold results.

Our study clearly demonstrates that low prediction quality often stems from an erroneous prediction of the elbow angle, with no major issue in VH-VL orientation prediction. Interestingly, there is some evidence in the literature that antibody elbow angles are influenced by the light chain class, and that Fabs with lambda chains have a wider range of angles (62). Moreover, various structural studies have reported that elbow angles are often altered by conformational changes that occur upon binding and give rise to significant differences between the elbow angles of the bound versus unbound Fabs (62, 81, 82). Notably, the nature of the antigen can affect the elbow bending angles within a given Fab, for example the elbow angle of anti-HIV-1 V3 Fab 2219 changes from 210.4 degrees upon binding to UG1033 peptide (PDB identifier 2B1A) to 229.4 degrees upon binding to MN peptide (PDB identifier 2B0S). Clearly, reliable prediction of the elbow angle in bound and unbound antibodies is a challenge that computational tools still have to address.

Docking by both ZDOCK and AlphaFold-Multimer predicted the structure of antibody:antigen complexes relatively poorly, with ZDOCK succeeding in a higher number of complexes. We note several limitations in our comparison. First, the input for the two alternatives is different, where the input for the viable docking option is the native structures of both the antigen and the antibody, while the AlphaFold prediction is based solely on the amino-acid sequences. Second, only 26 complexes were used for the comparison. It is important to note that all the complexes analyzed were released after the AlphaFold training cut-off date to avoid bias. Nevertheless, our study of 26 different complexes offers a sneak-peek at the current level of prediction accuracy expected of Ab-Ag complex structures.

The main conclusion of our study is that despite significant advancement in the ability to predict protein structures, the current methods are not very accurate at modeling the 3D structure of antibodies. This is predominantly due to low accuracy in prediction of the variable domains of the heavy chains, difficulty in predicting the CDR regions (in both heavy and light chains), and inability to model the elbow angle between the constant and variable domains correctly. Any methodology for predicting antibody structures should specifically address these hurdles. Future improvements in methodology should also improve computational-based epitope mapping algorithms.

## Acknowledgments

Israel Science Foundation (ISF) [2818/21] to T.P and [3136/22] to NTF; Binational Science Foundation (BSF) [01031771] to NTF; Edmond J. Safra Center for Bioinformatics at Tel Aviv University Fellowship to K.P. T.P.’s research is supported in part by the Edouard Seroussi Chair for Protein Nanobiotechnology, Tel Aviv University.

## Authors contribution

KP planned and performed the experiments, analysed data, prepared the figures, and wrote the manuscript together with NTF and TP. NTF and TP planned and supervised the experiments and data analyses, and wrote the manuscript together with KP.

## Inclusion and Ethics statement

All the authors meet the authorship criteria and gave their consent to be listed as authors on this manuscript.

