## Supplementary Figures for "Evaluation of the Ability of AlphaFold to Predict the Three-Dimensional Structures of Antibodies and Epitopes"

Supplementary Figure S1: Prediction quality evaluation by chain type and light chain class using the GDT_TS criterion.

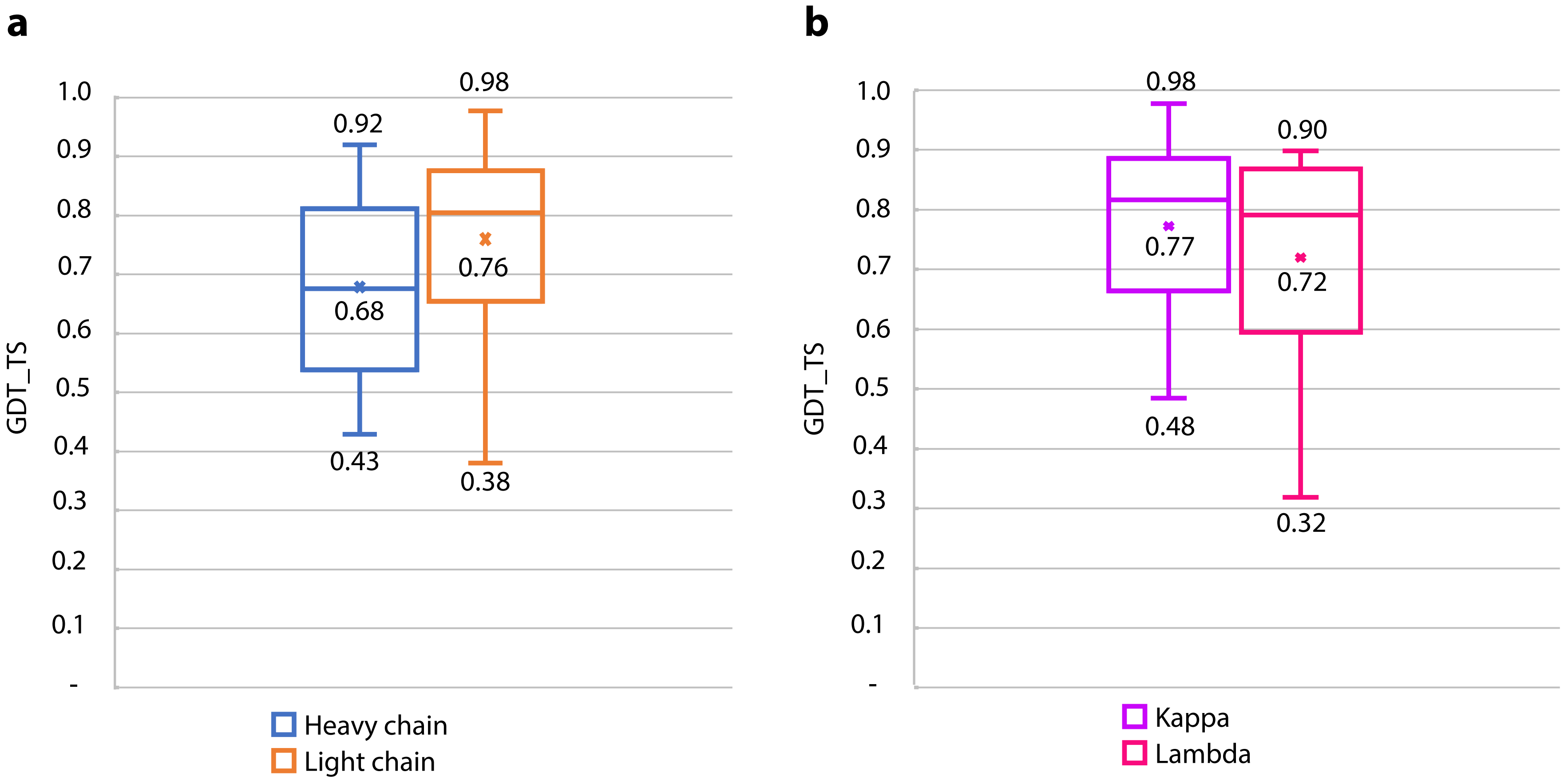


Boxplot of global distance test total score (GDT_TS) data displaying the similarity between superimposed native and predicted molecules for (a) light and heavy chains and (b) kappa and lambda chains.

Supplementary Figure S2: Prediction quality evaluation for different species and for different level of VDJ mutations.

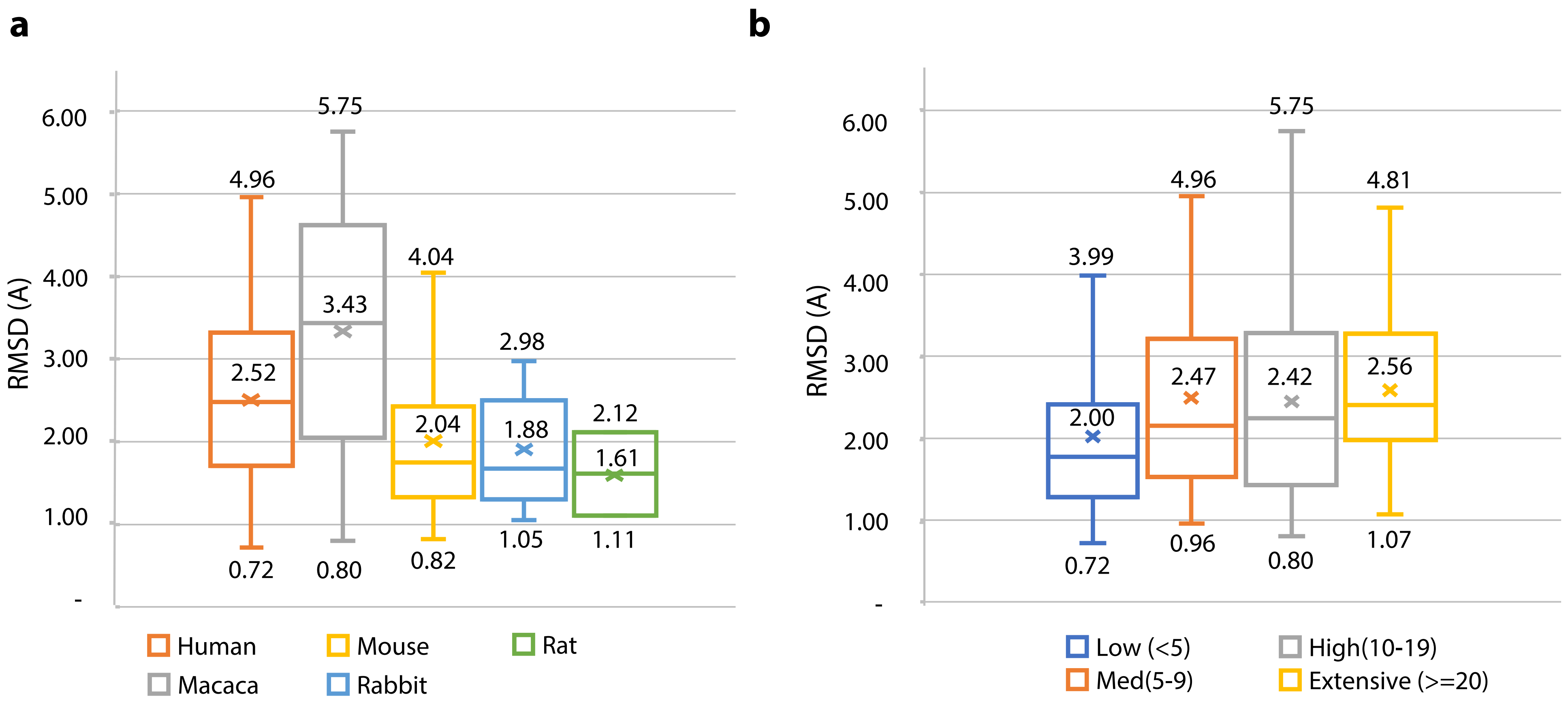


Boxplot of RMSD (Å ) data (a) for different species displaying the distance between superimposed native and predicted molecules; (b) for different level of VDJ mutations; Low level of mutations in the chain is defined as less than 5 amino acid substitutions, medium level - more or equal to 5 and less than 10, high level - more or equal to 10 and less than 20, and extensive level – more or equal to 20.
